# Population dynamics of underdominance gene drive systems in continuous space

**DOI:** 10.1101/449355

**Authors:** Jackson Champer, Joanna Zhao, Joanna Zhao, Samuel E. Champer, Jingxian Liu, Philipp W. Messer

## Abstract

Underdominance gene drive systems promise a mechanism for rapidly spreading payload alleles through a local population while otherwise remaining confined, unable to spread into neighboring populations due to their frequency-dependent dynamics. Such systems could provide a new tool in the fight against vector-borne diseases by disseminating transgenic payloads through vector populations. If local confinement can indeed be achieved, the decision-making process for the release of such constructs would likely be considerably simpler compared to other gene drive mechanisms such as CRISPR homing drives. So far, the confinement ability of underdominance systems has only been demonstrated in models of panmictic populations linked by migration. How such systems would behave in realistic populations where individuals move over continuous space remains largely unknown. Here, we study several underdominance systems in continuous-space population models and show that their dynamics are drastically altered from those in panmictic populations. Specifically, we find that all underdominance systems we studied can fail to persist in such environments, even after successful local establishment. At the same time, we find that a two-locus two-toxin-antitoxin system can still successfully invade neighboring populations in many scenarios even under weak migration. This suggests that the parameter space for underdominance systems to both establish in a given region and remain confined to that region would likely be highly limited. Overall, these results indicate that spatial context must be considered when assessing strategies for the deployment of underdominance systems.

## INTRODUCTION

Gene drives are genetic constructs designed to self-propagate through a population^1–3^. Various potential applications of these systems have been proposed, such as the dissemination of transgenes in disease-carrying insects that would render them unable to transmit diseases such as malaria or dengue^1–3^. One class of gene drives are underdominance systems, in which heterozygotes for the drive allele have lower fitness than homozygotes and wild-type individuals. This results in a frequency-dependent rate of spread of the drive allele and an invasion frequency threshold. When present below this frequency, the drive allele will tend to decrease in frequency over time and thus eventually be lost from the population, and when introduced above this frequency, the drive allele is expected to increase in frequency and usually go to fixation. Compared to other forms of gene drives, such as homing drives and *Medea*, underdominance systems typically require very large transgenic insect releases to successfully spread through a population, thereby requiring a substantially greater initial effort to establish the drive. However, the main purported advantage of underdominance systems is that they can successfully establish after a large-scale release and then remain locally confined to a given region. This property could ease regulatory and public approval requirements compared to other drive systems. A few distinctly different forms of underdominance systems have been demonstrated thus far. The earliest systems involved the formation of reciprocal chromosomal translocations (RCT)^4–7^. More recently, toxin-antitoxin systems at a single genomic locus (1L1T) have been developed^8^, as well as two-allele toxin-antitoxin systems at a single locus (1L2T) or two genetically unlinked loci (2L2T)^9^.

Several recent theoretical studies have examined the population dynamics of such underdominance systems in specific population models. The 1L1T system was found to stably persist without invasion into a neighboring population under a wide range of parameters and fitness values^10,11^. It was also found to be stable in connected panmictic population networks, unless the migration rate was very high^12^. Introduction threshold frequencies needed for successful establishment in a single panmictic population have been calculated for both the 1L2T and 2L2T systems^13^,14, and their capacity to invade neighboring regions has also been studied, which was shown to be possible only with the 2L2T system^14^. The 2L2T system was also assessed for different release strategies^15,16^, mechanisms of action^16,17^, payload fitness effects^17^, and combinations with other systems^18^. Its ability to establish in a population was found to be fairly robust to variation of these parameters.

While the behavior of underdominance systems is reasonably well understood in panmictic environments, it is less clear how they may act in spatially-explicit scenarios. Since underdominance drives rely on relative allele frequencies, their behavior may be significantly changed in spatially continuous populations, where local allele frequencies may be very different from the average frequency in the whole population. In fact, the tendency of underdominance systems to quickly fixate or be eliminated after release results in a tendency to form regions comprised of either underdominance or wild-type alleles, with potentially interesting interactions at the boundaries of these regions that could affect the overall population dynamics. Only one study to date has investigated the performance of underdominance gene drive systems in a spatially-explicit population model, and this study used a grid approximation for modeling the spatial dynamics and only examined the 2L2T system^15^. It was found in this study that the release threshold required for successful invasion varied based on the exact pattern of release in a simple spatially-explicit population (central vs patchy, small area vs. large area), and that this threshold was different from panmictic populations. That spatial structure could critically impact the behavior of an underdominance system is also suggested by modeling studies of *Wolbachia* systems, which share threshold-dependent dynamics with underdominance systems and have been shown to exhibit distinct dynamics in spatial models^19–28^.

Here, we conduct a comprehensive investigation of the population dynamics and general performance of each of the four underdominance system mentioned above in several types of spatially-continuous environments. We first examine the ability of these systems to establish locally, clearing a geographic region of wild-type individuals. We then study their ability to persist and remain in that region despite possible migration from adjacent wild-type populations. Finally, we study their ability to invade and increase in frequency in adjacent wild-type populations. We find that the spatial structure of a population has a profound impact on the fate of an underdominance system after it establishes, often preventing it from persisting successfully, or in the case of the 2L2T system, remaining confined to its initial release region.

## MODEL

### Underdominance systems

We study four different types of underdominance systems (Figure 1), spanning a variety of mechanisms and invasion threshold frequencies. Their dynamics are further tunable by the fitness costs associated with the drive mechanism itself and a potential payload allele. The following four underdominance systems are modeled:

1. *One-locus one-toxin-antitoxin (1L1T) system.* This system consists of a single drive allele, inserted into a single genomic locus (Figure 1A). The drive allele encodes an RNAi targeting a haploinsufficient gene (the “target”), located at a different locus than the drive allele (it does not matter whether these loci are genetically linked). In addition, the drive allele contains a functioning copy of the gene (the “rescue gene”), sufficiently recoded such that it is not itself prone to being targeted by the drive’s RNAi. In a homozygote for the drive allele, the target gene will be silenced, but since two copies of the rescue gene are present, the function of the haploinsufficient target will be completely restored. In heterozygotes, however, only one copy of the rescue gene is present, resulting in a lower fitness compared to either homozygote. Such a system is therefore expected to produce frequency-dependent invasion dynamics. In a panmictic population model, the invasion frequency threshold will be 50% if we assume that drive and wild-type homozygotes each have equal fitness (Figure 2). More generally, the dynamics and invasion threshold will be determined by the relative fitness values of the heterozygote and drive homozygote in relation to the wild-type homozygote. In our model, we assume that the drive allele contains a payload that reduces fitness by a factor *F* ≤ 1 in drive homozygotes relative to wild-type homozygotes. The silencing of the haploinsufficient gene in heterozygotes additionally reduces fitness by a factor of 0.26, a value that is inspired by a previous experimental demonstration of such a system^8^, where homozygotes for the drive had a fitness of *F* = 0.71 and heterozygotes had a fitness of 0.22. Assuming that the fitness factors are multiplicative, heterozygotes then have a relative fitness of 0.26√*F* in our model.
2. *Reciprocal chromosomal translocation (RCT) system.* This system consists of two drive alleles, generated by reciprocal translocation of two different chromosomes (Figure 1B). In such a system, the only three viable genotypes are homozygotes for the translocated alleles at both chromosomes, heterozygotes for the translocated alleles at both chromosomes, and homozygotes for the wild-type alleles at both chromosomes. The viable heterozygotes will then have at most 50% viable offspring, while mating pairs of homozygotes of either type will have normal numbers of offspring. Thus, heterozygotes are again at a disadvantage, similar as in the 1L1T system, leading to frequency-dependent dynamics with an invasion threshold of 50% if the drive homozygotes have equal fitness as wild-type homozygotes (Figure 2). We will assume more generally that one of the translocated chromosomes also carries a payload gene, and that homozygotes for both translocations have a relative fitness *F* compared to homozygotes for the wild-type alleles at both chromosomes. The viable heterozygotes then have a relative fitness of √*F*.
3. *One-locus two-toxin-antitoxin (1L2T) system.* This system requires two different drive alleles at a single genomic locus, each encoding a different toxin, while at the same time carrying an RNAi (the “antidote”) that silences the toxin of the other drive allele (Figure 1C). In this case, the only viable genotypes are heterozygotes for the two drive alleles and homozygotes for the wild-type allele. Such a system would exhibit frequency-dependent dynamics with a comparatively high invasion threshold of 67% of two-drive allele heterozygotes, assuming that they have equal fitness as the wild-type homozygote (Figure 2). To model possible additional fitness costs, including those of a payload gene (which could be located inside one or both drive alleles), we assume that these heterozygotes have a relative fitness *F* compared to wild-type homozygotes.
4. *Two-locus two-toxin-antitoxin (2L2T) system.* Our final system is a variation of the 1L2T system in which the two drive alleles are inserted into two genetically unlinked genomic loci (Figure 1D). We further assume that the antidote from one copy of a drive allele can completely silence two toxin copies from the other drive allele. In that case, the only non-viable offspring would be those individuals that inherited either one or two copies of any single drive allele while not inheriting any copy of the other allele. In the absence of additional fitness costs, this would result in a comparatively low invasion threshold of 27% of drive allele homozygotes at both loci (Figure 2). We assume that one of the drive alleles could also contain a payload gene with a relative fitness *F* in homozygotes and √*F* in heterozygotes compared with wild-type individuals, while the drive allele at the other locus carries no payload or additional fitness cost. One interesting feature of this model is that in case of *F* < 1, the drive alleles do not actually go to fixation but reach an equilibrium with wild-type alleles (Figure 2).

**Figure 1.**
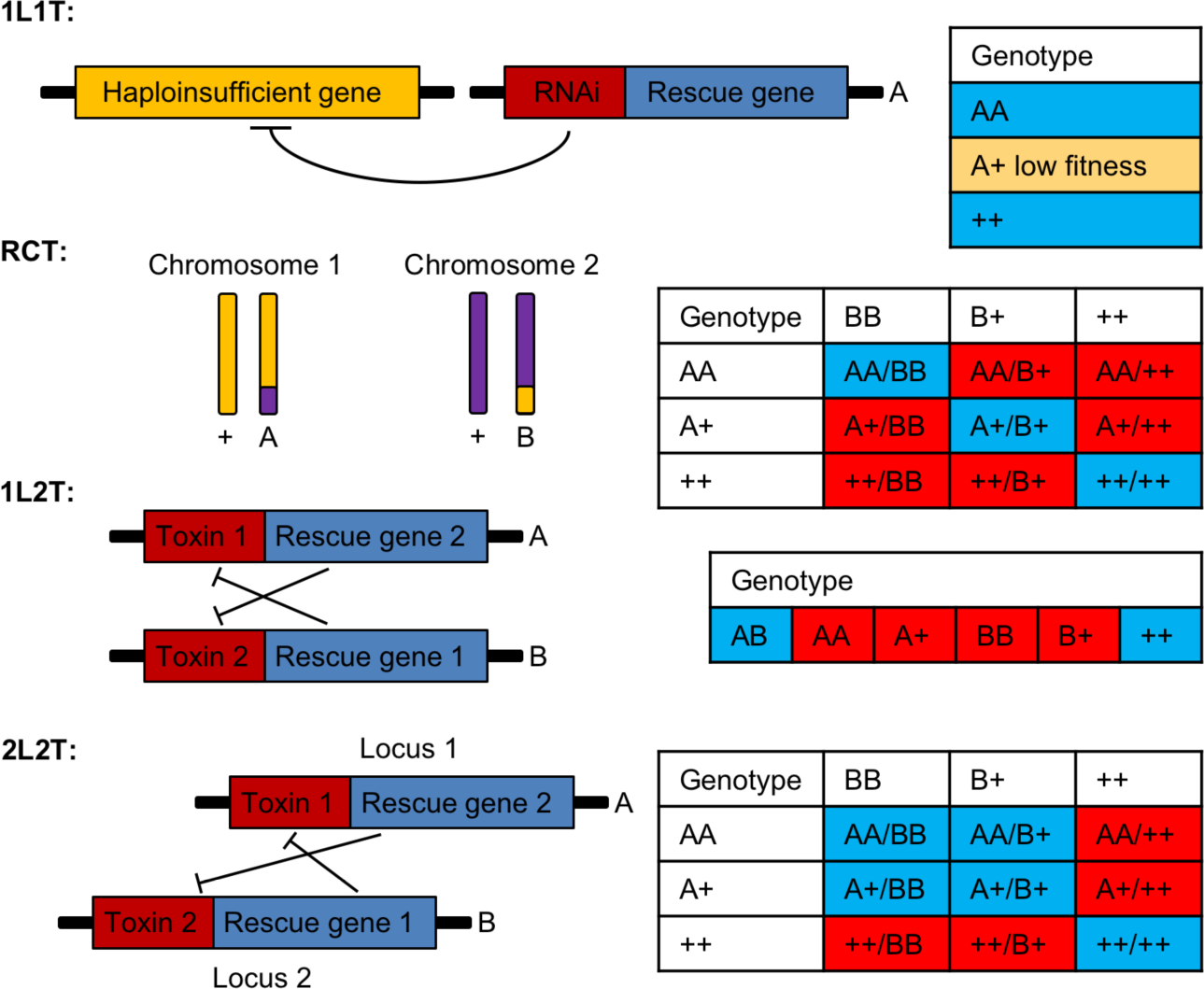
Underdominance systems. The figure illustrates the mechanisms of action and viable genotypes for each of the four drives (blue – viable, red – not viable). The 1L1T system works by targeting a haploinsufficient gene with RNAi. The RCT system involves translocations of two different chromosomes. The 1L2T system has two drive alleles, each containing a toxin that is compensated by a rescue gene in the other allele. Both alleles are located at the same locus and a single copy of a rescue gene can fully compensate two copies of the toxin. The 2L2T system is a variation of the 1L1T system where the two drive alleles are located at different loci.

**Figure 2.**
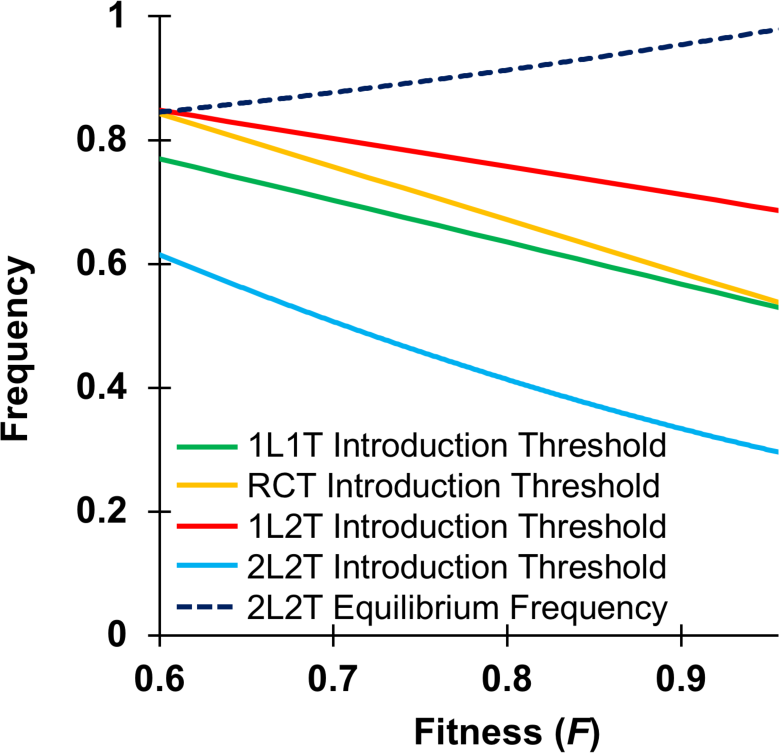
Threshold frequencies of underdominance systems. In a deterministic Wright-Fisher model of a single panmictic population, the invasion threshold frequencies of the four drive systems vary as a function of the fitness parameter *F*. These threshold frequencies represent unstable equilibria below which the drive is removed and above which the drive increases in frequency (either to fixation or to a high equilibrium frequency for the 2L2T drive, shown by the dashed black line).

We note that the experimental demonstrations of both the 1L2T and 2L2T systems are based on maternally-active *Medea* elements^9^, yet this system has proven difficult to transfer from *Drosophila* to other relevant target organisms. Thus, for our study we chose to model conceptually similar but more general underdominance systems in which the toxin and antitoxin effects are based on the genotype of any individual, regardless of parent.

### Basic panmictic population model

We implemented each of the four underdominance systems in an individual-based, forward-in-time population genetic simulation framework, using the SLiM software version 3.1^29,30^. Here we will first outline the general life cycle of these simulations for the basic panmictic population model. The extension to continuous space will be discussed below.

Our basic panmictic model simulates a population of sexually-reproducing diploid hermaphrodites. The life cycle is based broadly on a Wright-Fisher-type model, where individuals evolve over discrete generation during which they reproduce and then die. To obtain the individuals of the next generation, we first calculate how many such individuals will be generated. In our model, this is determined by the number of individuals in the previous generation and the carrying capacity of the population. Specifically, if *N* individuals were present in generation *t*, we will generate 10*N*/(1+9*N*/*K*) children in generation *t*+1, where *K* is the carrying capacity. This model produces logistic dynamics and was selected to smoothly but quickly restore the population to carrying capacity after perturbation (e.g. after the initial release of drive-carrying individuals into a wild-type population). The population size would be expected to approach *K* under this model if all children were viable. However, for our underdominance systems there can also be non-viable offspring, so the actual population size is often below carrying capacity.

The next step is then to generate these individuals. For each one, we first draw a random individual from generation *t* as its first parent. The probability to be drawn is proportional to each individual’s fitness. A second parent is drawn accordingly. The child’s genotype is then determined by Mendelian inheritance. After all children have been generated this way, we remove the non-viable children from the population. The remaining individuals then serve as the parents for the individuals of generation *t*+2.

In our analyses of the behavior of the different underdominance systems in the panmictic population model, we will focus on a specific scenario of two subpopulations linked by migration at rate *m*, specifying the probability that a child created in one population has its parents drawn from the other population. The specific setup of this scenario and the parameters used will be detailed in the Analysis section.

### Extension to continuous space

To modify the general panmictic model to 2D continuous space, four fundamental aspects must be modified. First, each individual now has properties representing their *x* and *y* coordinates in space. These coordinates are set when a new child is generated, by displacing its position from that of its first parent into a random direction and by a distance drawn from an exponential distribution with mean *m* (the behavior at the boundaries of the space will be described below).

Next, we define how spatiality impacts the mating behavior of individuals. In the panmictic model, mates could be picked among all existing individuals. In the spatial model, we assume that only mates within a certain radius of the first parent can be chosen. For simplicity, we further assume that this mating radius is equal to the average dispersal distance at which new offspring are placed from the first parent, which we simply term the “migration rate” (*m*) of the spatial model. The probability of potential mates to be picked is again proportional to their fitness. If no possible mate is present within the specified area, the first parent is redrawn until a child has been successfully generated.

The third aspect we change for the spatial model is that we adopt a mechanism of local density regulation. This will be critical for modeling realistic spatial dynamics, which should often be determined by local competition. Such density regulation will also prevent unrealistic agglomeration of individuals in small areas. We implement local density regulation by defining a carrying density r*_K_* = *K*/*A*, where *K* is the global carrying capacity and *A* is the total area in which the population lives. For every individual in the population, we calculate the local density r*_i_* of individuals present within a circle of radius *r* centered around its location. The fitness of the individual is then rescaled using the same logistic form we used for the global carrying capacity in the panmictic scenario but using densities instead: w*_i_*’ = w*_i_* × 10/(1+9r*_i_*/r*_K_*). Individuals located in more dense areas will thereby tend to have lower fitness, and thus fewer children, which will attenuate regional density fluctuations.

Finally, we need to specify how many children will actually be generated in the next generation. In the panmictic model, this was determined by the global carrying capacity. However, if we were to adopt the same approach in the spatial model, this would often result in biologically unrealistic behavior due to artifacts resulting from Wright-Fisher assumptions. For example, if density was very high in a tiny region containing most individuals with only very few individuals elsewhere, the Wright-Fisher model would assign far more offspring to these few individuals than might be biologically possible. To address this, we set the overall number of individuals present in the next generation to be the sum of the net fitness of individuals in the previous generation. For the RCT, 1L2T, and 2L2T systems, where a fraction of offspring is expected to be non-viable, we further assume that some individuals only contributed a fraction of their net fitness to the count of offspring in the next generation (details are described in the SLiM configuration files for these models).

These complexities illustrate how the Wright-Fisher model approaches its limits for modeling spatial populations. More realistic models would likely depart from this framework and instead model mating events and litter sizes explicitly, such that the population size in the next generation would emerge naturally from these events. While this is possible in the latest versions of SLiM, we used a Wright-Fisher framework for our spatial models to ensure comparability with the panmictic scenario.

In our analyses of the behavior of the different underdominance systems in the spatial population models, we will study four specific scenarios defined by the geometry of the spatial area and the initial distribution of drive individuals. These scenarios will be described in detail in the Analysis section.

### Data generation

Simulations were run on the high-performance computing cluster of the Department of Biological Statistics and Computational Biology at Cornell University. Data processing, analyses, and figure preparation were performed in R. All SLiM configuration files for the implementation of the different underdominance strategies and study scenarios and all simulation data are available on GitHub (https://github.com/MesserLab/UnderdominanceGeneDriveSystems).

## ANALYSIS

### Panmictic release scenario

The panmictic release scenario (Figure 3A) models two panmictic populations connected by migration and serves to create the baseline for comparison to our spatial scenarios. We assumed that both populations have equal carrying capacity (*K* = 10,000) and are linked by symmetric migration at rate *m*. The scenario was initialized by first introducing 10,000 wild-type individuals into each population. Homozygotes for the gene drive alleles (or heterozygotes for the two different drive alleles in the 1L2T system) were then added to the existing individuals in the first population so that they represented 80% of that population at the time of release. This frequency is usually well above the expected invasion thresholds for each of the four underdominance systems, even for strong fitness costs.

**Figure 3.**
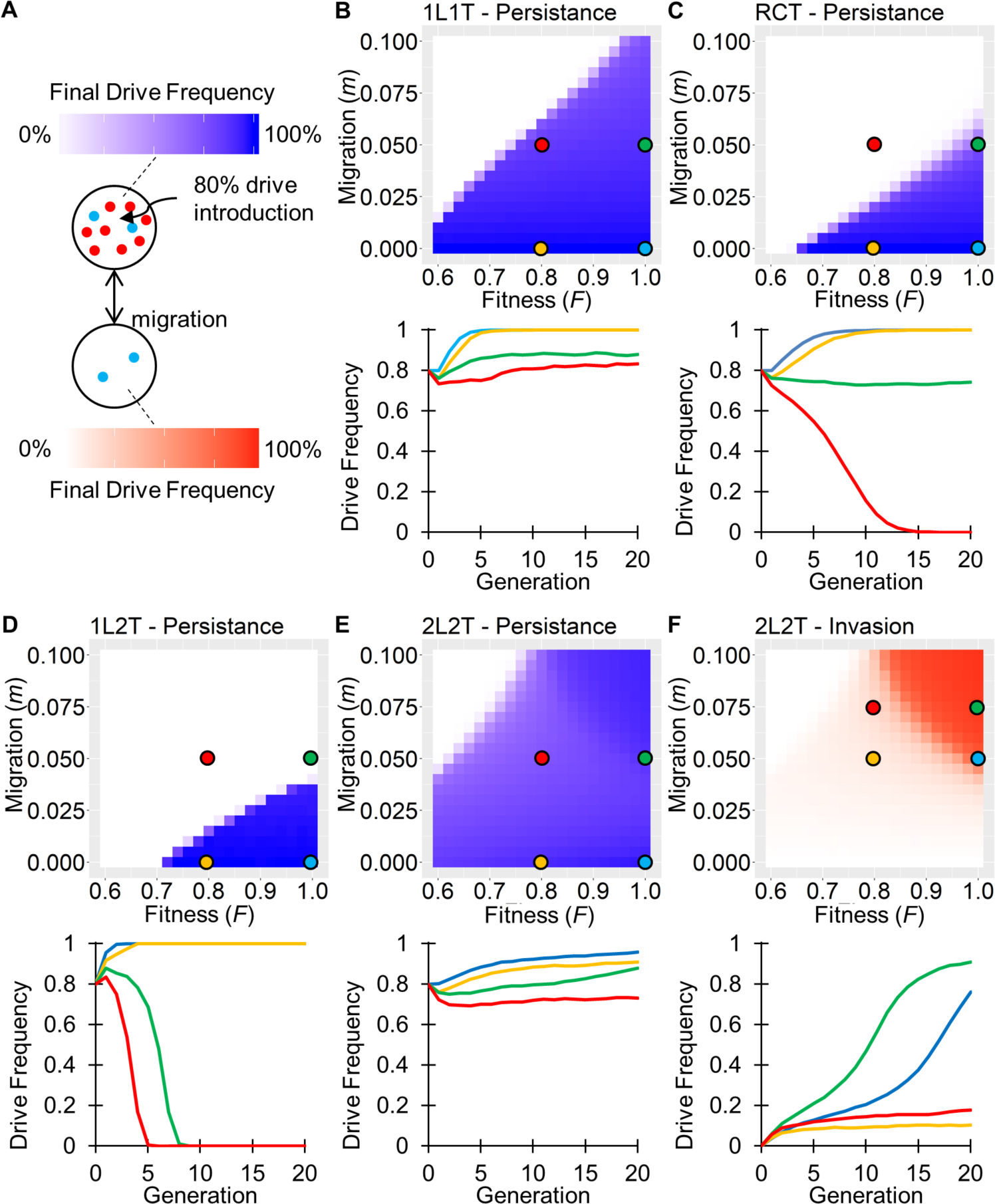
Panmictic release scenario. (A) Two panmictic populations of constant size are linked by symmetric migration. Drive individuals are introduced to the first population such that they constitute 80% frequency. (B)-(E) Ability of the four underdominance systems to persist in the release population after 20 generations. (F) Ability of the 2L2T system to invade the adjoining population after 20 generations. Each data point in the heatmaps was obtained by averaging the final drive allele frequency over 10 independent simulation runs for the given parameter setting. The graphs below each heatmap show drive frequency trajectories from individual simulations for parameter settings marked by the circles on the heatmaps with corresponding colors. Persistence and invasion heatmaps for all four systems and comparison to a non-driving allele are shown in Figure S1.

We analyzed the population dynamics of the four underdominance systems over a grid of parameter values in which we varied the fitness parameter *F* of each system from 0.6 to 1.0 and the migration rate *m* from 0.0 to 0.1. To assess the ability of the drive to persist in the release population, we recorded the final drive frequencies in the release population after 20 generations. To assess the ability of the drive system to invade a separate population of wild-type individuals, we recorded the final drive frequency in the other population.

Consistent with previous studies, we found that as long as fitness costs and migration rates were sufficiently low, all four underdominance systems were able to successfully establish (i.e. initially increase in frequency in the release population to nearly 100%, or the equilibrium frequency for the 2L2T system), and then persist in the release population over the course of 20 generations (Figure 3B-E). The RCT system had more trouble persisting in the presence of high migration than the 1L1T system, presumably due to its lower ability to remove wild-type individuals. As expected, the 1L2T system was the most sensitive to migration of all the underdominance systems because of its high invasion threshold, where even low immigration rates of wild-type individuals sufficed to push the drive below its threshold in the release population. However, the 1L2T system did establish more quickly in the release population than the other systems. The 2L2T system established the most slowly, but it was the most persistent in the face of migration, even when fitness costs were high. Moreover, when fitness costs were sufficiently small and migration sufficiently high, this system was able to invade the adjoining population (Figure 3F). Overall, these results largely recapitulated those from earlier studies^10–18^.

### Spatial linear scenario

In this scenario, we study a spatially continuous population to explore the behavior at the boundary between a wild-type region and adjoining region in which an underdominance system has established (Figure 4A). Specifically, we modeled a square arena of global carrying capacity *K* = 10,000 with edge-length defined as 1 length unit. The total area is therefore *A* = 1 and the carrying density is r*_K_* = *K*. We further set *r* = 0.02 (i.e., local population densities were estimated over a circle of radius 2% of the edge-length of the arena). This value was chosen so that the expected number of individuals in such a circle would be sufficiently high for a small variation in number to not have a large impact on fitness. We used reprising boundary conditions for the arena, meaning that if a new child was assigned coordinates that would have placed it outside of the boundaries, its position was redrawn until it fell inside the arena. The scenario was initialized by randomly placing 5,000 drive homozygotes into the left half of the arena and 5,000 wild-type homozygotes into the right half. After 60 generations, we recorded drive allele frequencies in the left half to assess persistence and the right half to assess invasion. From these data, heatmaps were generated similarly to the panmictic release scenario, except that for all spatial scenarios, we only studied migration rates *m* ≥ 0.02 to avoid excessive clustering of individuals.

**Figure 4.**
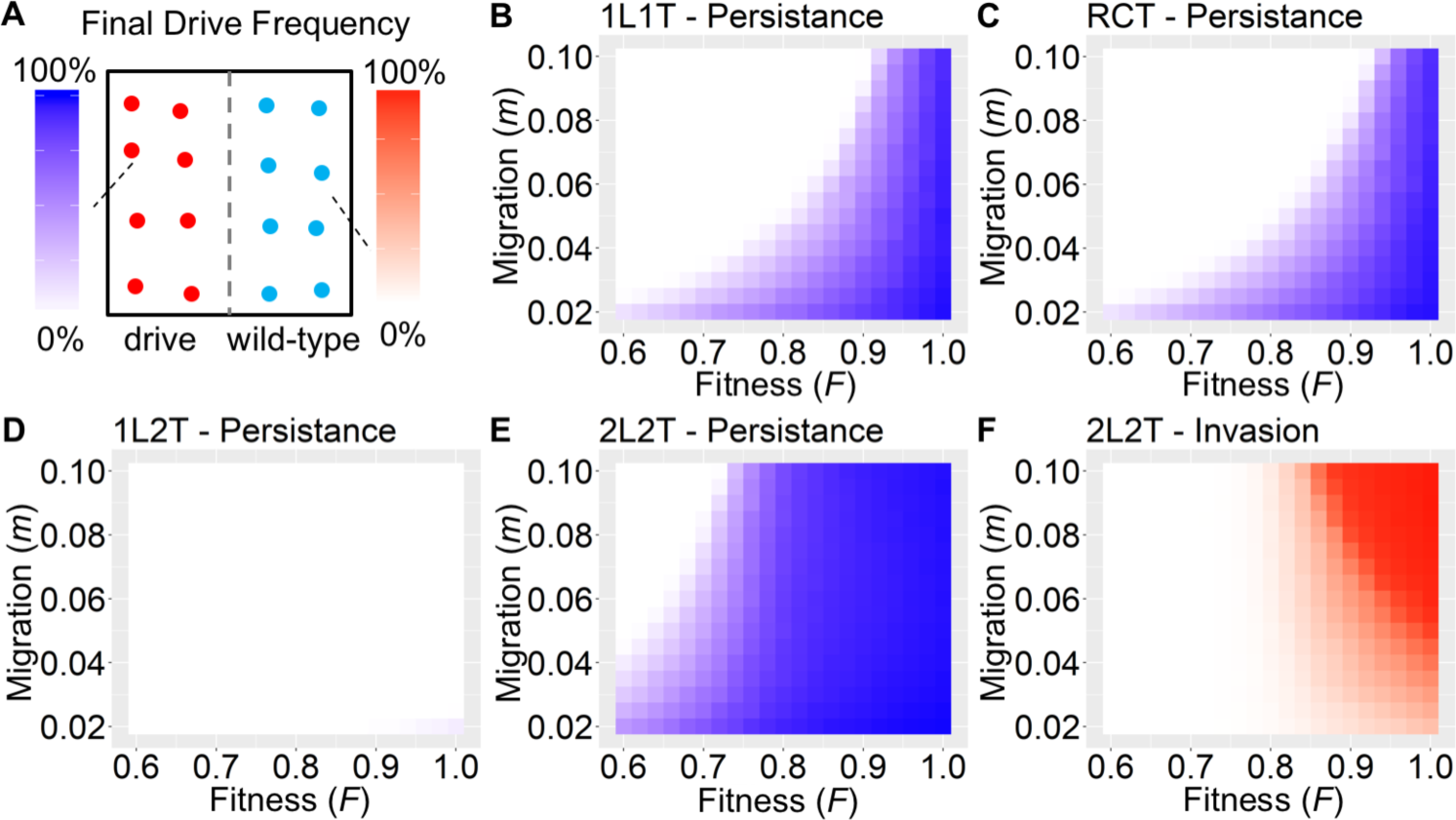
Spatial linear scenario. (A) The square arena is initialized with drive individuals in the left half and wild-type individuals in the right half. (B)-(E) Ability of the four underdominance systems to persist in the left half after 60 generations. (F) Ability of the 2L2T system to invade the right half after 60 generations. Persistence and invasion heatmaps for all four systems and comparison to a non-driving allele are shown in Figure S2.

We first checked that when a non-driving allele was introduced instead of an underdominance system, it indeed diffused quickly throughout the population (Figure S2). By contrast, all underdominance systems were able to maintain a distinct boundary between the region with drive individuals and the region with wild-type individuals, with the sharpness of the boundary depending on the migration rate and type of drive based on its ability to suppress heterozygotes (1L2T, 1L1T, RCT, and 2L2T in declining order of sharpness). The 1L1T and RCT systems were able to maintain the boundary at nearly the same location over the whole 60 generations for *F* = 0, but with increasing fitness costs, the boundary started receding into the left half of the arena (Figure 4B-C). The 1L2T system was rapidly eliminated from the entire arena, even with no fitness costs and low migration (Figure 4D). The 2L2T system was capable of persisting in its release region even with moderate fitness costs (Figure 4E). For low fitness costs, it could also successfully invade the wild-type region (Figure 4F). Higher migration generally accelerated the observed behavior in all these cases.

### Spatial circular release scenarios

This scenario models a square arena, similar to the spatial linear model, but drive individuals are released in a circle located in the center of the arena (Figure 5A). Such a scenario should be more representative of realistic releases and allows us to assess the effect of a curved boundary, as compared to the linear boundary of the previous scenario. The model was initialized by randomly placing 10,000 wild-type individuals uniformly across the entire arena, which we modeled with periodic (toroidal) boundaries in this scenario. The population was then allowed to equilibrate for 10 generations, at which point drive individuals were released into a central circle of radius 0.25 in sufficient quantity to represent approximately 80% of the individuals in this region. For all four underdominance systems, this release size assured initial establishment in the release circle under most of the parameters studied. After 30 generations, we recorded drive allele frequencies inside the release circle to assess persistence, and outside of the release circle to assess invasion.

All four underdominance systems were generally less effective at persisting in this circular release scenario than in the spatial linear scenario, which is a general consequence of the different geometries of these scenarios. In the linear scenario, an individual located at the boundary between the drive and wild-type areas would observe a drive allele frequency of ~50% when estimated over a small circle around its location, which is exactly the threshold frequency for the 1L1T and RCT systems when *F* = 0, resulting in a stable boundary. In the circular scenario, however, drive individuals encounter wild-type individuals at a concave boundary, while wild-type individuals encounter drive individuals at a convex boundary. An individual located at the boundary will thus observe a local drive allele frequency of <50% in a small circle around its location, which is below the threshold frequency of the 1L1T and RCT systems. Indeed, we observed that even in the absence of fitness costs, the 1L1T and RCT systems both receded in this scenario across the entire parameter range and were usually completely eliminated after 30 generations (Figure 5B-C). Increasing fitness costs and migration further accelerated this process. The 1L2T system was generally the fastest to recede among all systems (Figure 5D). The 2L2T system was able to persist for low fitness costs and low migration (Figure 5E) and could even begin to invade the surrounding area at somewhat higher migration (Figure 5F). Yet, the parameter range under which this was observed was much more limited than in the linear scenario. These patterns indicate that even if a 50% threshold drive has the same fitness as wild-type alleles, it will still recede if at a geometrical disadvantage, which restricts the possible release profile for successful use of an underdominance drive.

**Figure 5.**
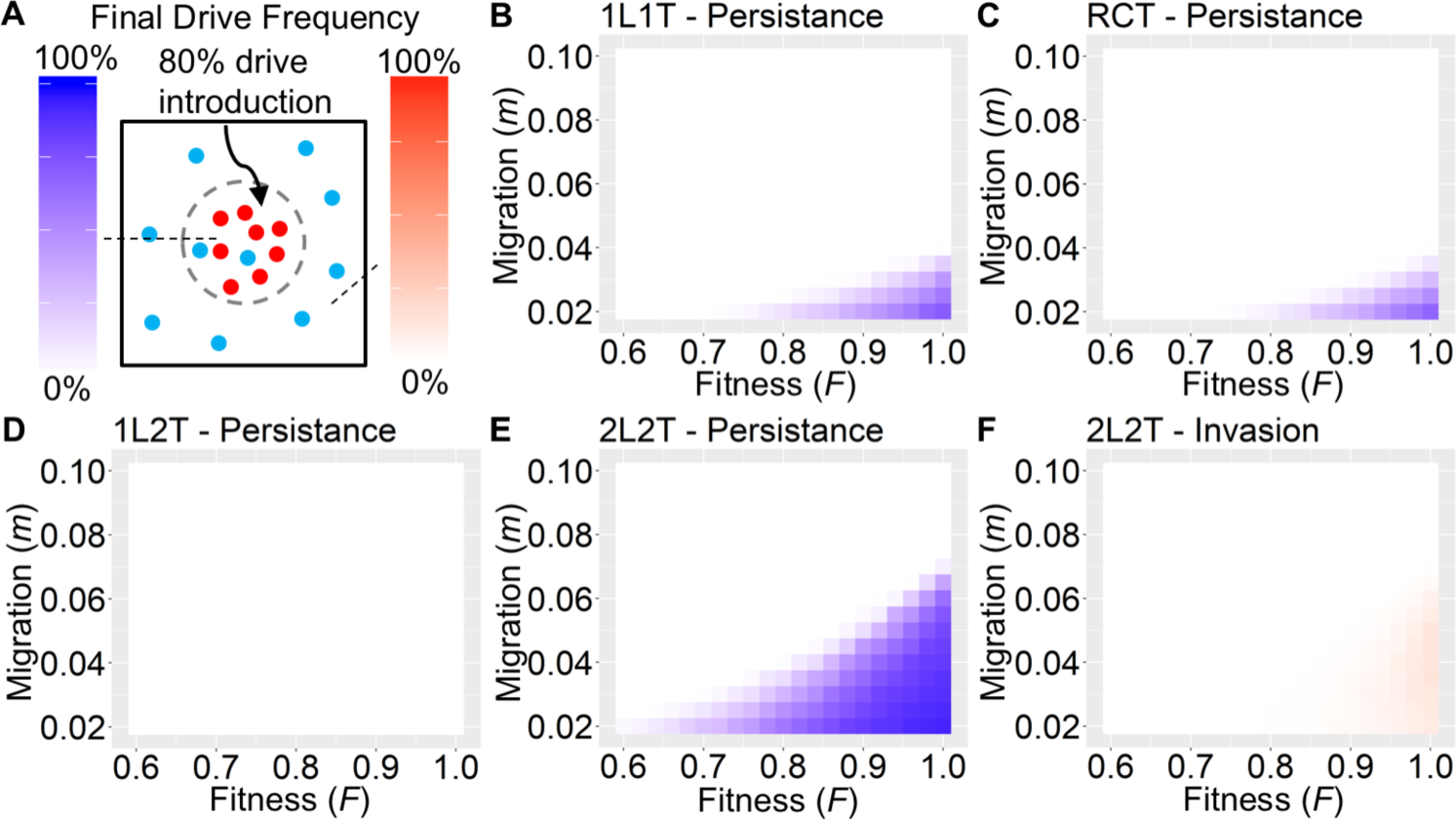
Spatial circular release scenarios. (A) Square arena with drive individuals released in a central circle. (B)-(E) Ability of the four underdominance systems to persist inside the area of the initial release circle after 30 generations. (F) Ability of the 2L2T system to invade the area outside of the initial release circle after 30 generations. Persistence and invasion heatmaps for all four systems and comparison to a non-driving allele are shown in Figure S3.

### Spatial high-density circular release scenario

Certain applications of underdominance systems may involve the release into an area of high population density, such as a city or wetland region, which is surrounded by a region of lower density. To model such a scenario, we modified the circular release scenario described above so that the carrying density inside the central release circle could be increased by a factor *x*, while keeping the carrying density outside of the circle the same as before (Figure 6A). For a given value of *x*, the release number was then adjusted accordingly such that drive individuals still initially constituted 80% of individuals inside the release circle. Persistence and invasion ability were again measured after 30 generations.

When we doubled the carrying density of the release circle compared to that of the surrounding region (*x* = 2), both the 1L1T and RCT systems were now able to persist in the release area for low fitness costs and migration (Figure 6B-C). At equilibrium, the boundary typically extended somewhat beyond the initial release circle. As carrying density in the center was further increased, the 1L1T and RCT systems were even able to invade their surrounding area for low fitness costs and high migration, presumably due to high diffusion from the circle into the outside areas (Figure S6B). However, in a larger arena, this apparent invasion would presumably take the form of an equilibrium with surrounding wild-type individuals, and the drive would still remain confined to the area surrounding the region of higher density. Thus, for the 1L1T and RCT drives, a successful but confined release could potentially take place in a region of higher density if the release encompasses nearly the entire region. The 1L2T drive was unable to persist over most of the examined parameter space, except for very high carrying densities in the release circle and low fitness costs (Figure 6D). With 2-fold higher central population density, the 2L2T drive could persist over a wide range of parameters (Figure 6E) and quickly began to invade the surrounding area unless it had a high fitness cost (Figure 6F).

**Figure 6.**
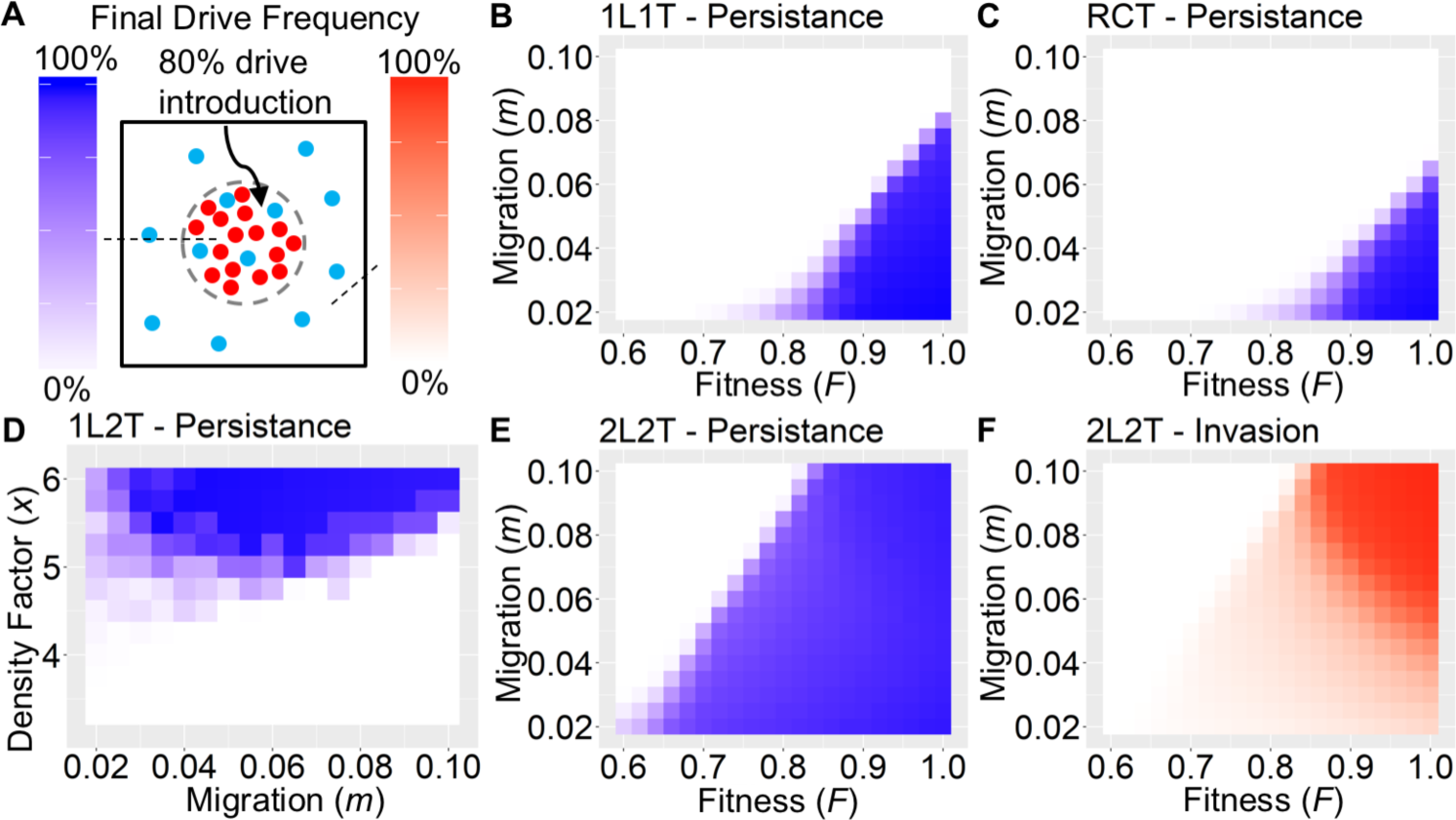
Spatial high-density circle release scenario. (A) Rectangular arena with drive individuals released in a central circle of *x*-fold higher carrying density compared to the surrounding area. (B) Persistence of the 1L1T system for *x* = 2. (C) Persistence of the RCT system for *x* = 2. Persistence of the 1L2T system as a function of *x* and migration for *F* = 0. Persistence of the 1L2T system for *x* = 2. (F) invasion of the 2L2T drive for *x* = 2. Figures S4-S6 show additional persistence and invasion heatmaps for all four systems and additional parameter variations.

### Spatial migration corridor scenario

In the panmictic release scenario we modeled two panmictic populations connected by migration, where an underdominance systems could establish in one of the populations while the other remained wild type. To study the extension of such a model to continuous space, we considered two circular populations, each with radius 0.4, and connected by a “migration corridor” of length 0.4 and variable width *w*, assuming reprising boundary conditions (Figure 7A). The total area of the arena is therefore ≈ 2π*0.4^2^+0.4*w*. Assuming a uniform carrying density of r*_K_* = 10,000, this yields an expected population size of ≈ 10,000+4,000*w* individuals. We initialized the model with a randomly distributed population of this size. All individuals in the left circle and left half of the migration corridor were drive individuals, while all individuals in the right half were wild type. After 100 generations, we recorded drive allele frequencies in the left half of the arena to assess persistence, and in the right half to assess invasion.

For a corridor width of *w* = 0.1, both the 1L1T and RCT systems were able to persist with even moderate fitness costs (Figure 7B-C), receding only inside the corridor region. The 1L2T system could persist successfully only in the absence of fitness costs and when the corridor width was small (Figure 7D). Otherwise it quickly receded. Interestingly, a moderate rather than low level of migration was actually beneficial toward the persistence of these drives, presumably due to the specific geometry at the interface between the circle and the corridor, where wild-type individuals trying to immigrate into the left circle would have a hard time reaching high-enough local frequency in the presence of strong diffusion. The 2L2T system, on the other hand, was able to persist in the left half even with high fitness costs (Figure 7E), and it could also invade the right half when corridor width and migration was high and fitness costs were small (Figure 7F). For intermediate corridor width, a more moderate level of migration was optimal for invasion. Thus, under certain conditions, underdominance systems can persist in one region while failing to invade a connected region.

**Figure 7.**
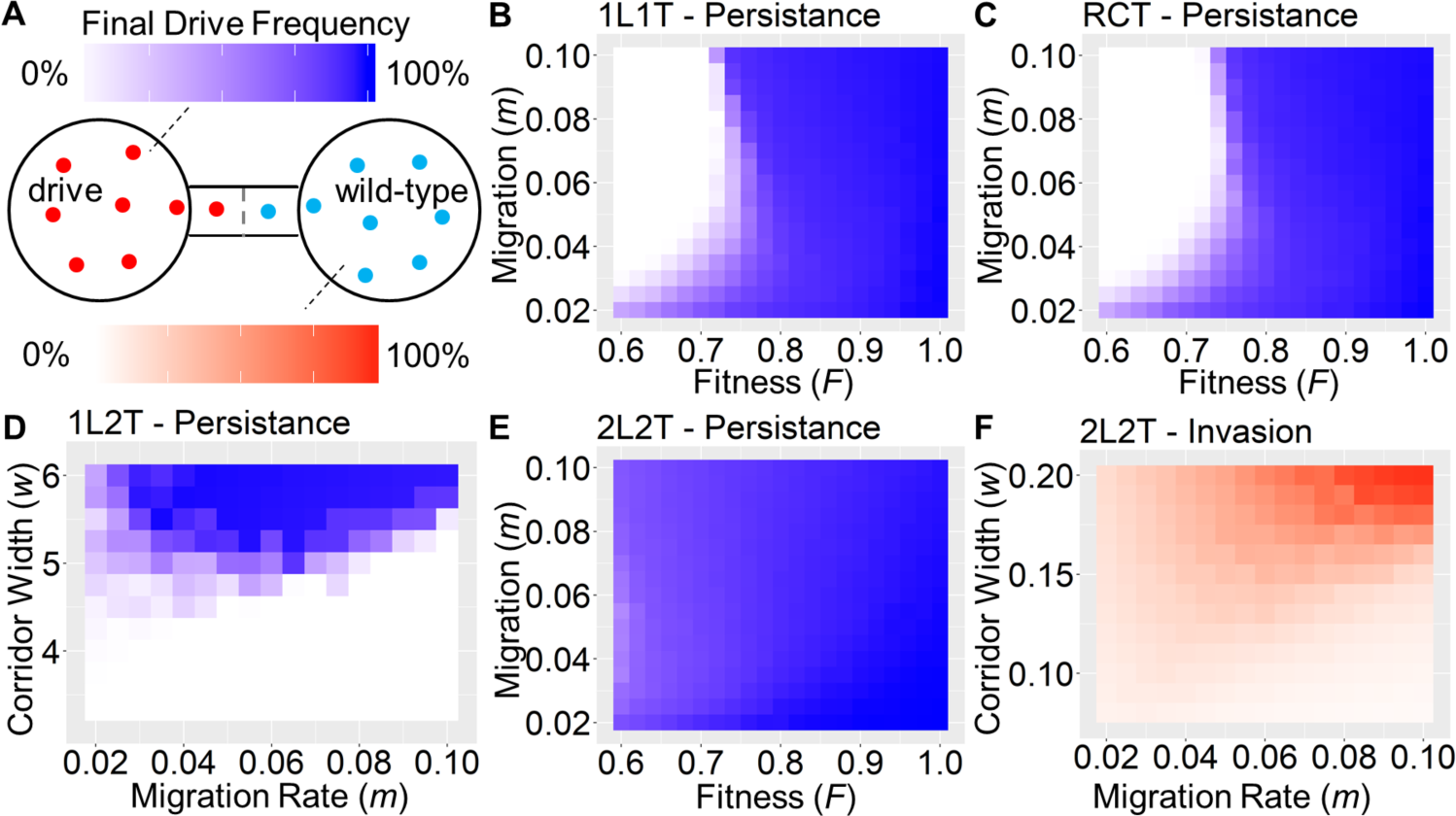
Spatial migration corridor scenario. (A) Two circular areas are connected by a migration corridor, with drive individuals initially present in the left half of the arena and wild-type individuals in the right half. (B) Persistence of the 1L1T system for *w* = 0.1. (C) Persistence of the RCT system for *w* = 0.1. (D) Persistence of the 1L2T system as a function of *w* and migration for *F* = 0. (E) Persistence of the 2L2T system for *w* = 0.1. (F) Persistence of the 2L2T system as a function of *w* and migration for *F* = 0. Figures S7-S9 show additional persistence and invasion heatmaps for all four systems and additional parameter variations.

## DISCUSSION

The key appeal of underdominance systems lies in the promise of a drive mechanism that could remain locally confined due to frequency-dependent threshold dynamics. In a panmictic population, the frequency of a drive allele is a well-defined parameter, making it straightforward to predict whether a given underdominance system should increase or decreased in frequency. After successful establishment in one population, the drive can be prevented from invading other populations when migration is too weak to push it above its threshold. However, most real-world populations are inherently spatial, with individuals moving over a continuous landscape. Here the frequency of a drive allele becomes a local parameter that can vary over the landscape, and the effects of migration can be very different than in a panmictic model. For instance, in a spatial population, migration often proceeds through spatial corridors. In this case, there will be areas with substantially higher local immigrant frequencies that could serve as “seeds” for invasion even if migration is comparatively low overall.

In this study, we demonstrated that the dynamics of underdominance systems are indeed substantially different in such spatially-continuous populations than in panmictic populations. This is true for their ability to persist in a release area, as well as their ability to subsequently invade a neighboring population. While all four underdominance systems we studied (RCT, 1L1T, 1L2T, and 2L2T) were able to establish and persist in a panmictic population when released at sufficient frequency, the scenarios in which they were able to persist in a spatial population were much more limited. For example, we found that only systems with an invasion threshold of 50% or lower in a panmictic population were able to maintain a stable boundary in the spatial linear scenario, while all other systems were steadily rolled back. This means that only the 2L2T system, as well as the RCT and 1L1T systems with no fitness costs, would be able to persist in such a scenario. In the circular release scenarios, even the 50%-threshold drives were quickly replaced by wild-type individuals, and only the low-threshold 2L2T system (with low fitness costs) was in fact capable of persisting. When sufficiently increasing the carrying density of the release area compared to the surrounding area, all four systems were again able to persist inside the release circle.

These results can be understood qualitatively when considering the local neighborhood located at the boundary between the drive and wild-type regions (where we define neighborhood by the average dispersal/mating distance of individuals). For an individual at such a location, drive alleles will tend to be at a frequency of 50% if the boundary is straight (as in the linear scenario) and at a lower frequency when the boundary is concave (as in the circular scenario), assuming that the density of individuals is equal in all areas. For the boundary to be stable, the underdominance system needs to have an invasion threshold close to this frequency. If its threshold is lower, the boundary will recede. Increasing the density of individuals in the drive area will increase the local drive allele frequency at the boundary, explaining why the underdominance systems were more effective in this case. On the other hand, decreasing the drive density due to fitness costs will have the opposite effect, in addition to direct effects of the fitness cost on the drive threshold.

The low-threshold 2L2T system was the most capable of establishing and persisting, but this also made it quite capable of invading neighboring regions. For sufficiently high migration, this system was able to quickly advance its boundary in the linear scenario and was able to invade even in the circular release scenario. It was also often able to invade through a narrow migration corridor. Therefore, the 2L2T system must be considered invasive in the absence of moderate to high fitness costs for a wide range of possible scenarios. This is in marked contrast to panmictic scenarios, where this systems was not found to be invasive for low migration rates^14^.

When comparing the relative merits of the four different underdominance systems in spatial scenarios, several patterns emerge. While the 1L2T can establish in a local region the most quickly, it is highly vulnerable to all but the lowest levels of migration, and even a narrow migration corridor can result in successful invasion of wild-type individuals. Thus, this system would likely be of use only on small islands or in otherwise highly isolated regions with low levels of migration, given that wild-type individuals would quickly be able to reclaim the area even from a small initial refuge. The 2L2T system establishes the most slowly but can persist in numerous scenarios due to its low threshold. Problematically, this also made it the most invasive. The intermediate threshold 1L1T and RCT systems can establish quickly, but have trouble persisting in several realistic scenarios. Comparing the two systems, RCT has lower ability to suppress wild-type alleles, making it more vulnerable to migration than 1L1T, particularly during initial establishment. On the other hand, RCT alleles are likely less vulnerable to genomic instability causing the separation of toxin and antitoxin components. This advantage may be of low importance compared to other underdominance systems, particularly the 1L1T RPM-Drive system, which was recently shown to be highly stable for an extended period of time^31^. RCT systems additionally would suffer from reduced levels of crossovers^32^, reducing the introgression of higher fitness wild-type chromosomes, and effectively adding a fitness cost after establishment of the drives that we did not model. For other systems, this reduction in fitness from laboratory genetic backgrounds compared to wild-type individuals could be overcome by crossing of wild-type individuals to drive individuals prior to generating homozygous stocks for release. Without such a method, all underdominance systems would likely suffer substantial fitness costs compared to wild-type individuals, resulting in higher release thresholds and reduced chances of successful persistence.

Our results suggest two possible strategies for increasing the chance that an underdominance drive can successfully establish and persist in a spatial population. The first strategy is to ensure that the release covers the entire area of the target population at a level well above the threshold for a high-threshold drive. For a medium-threshold drive, coverage over at least most of the area is necessary, so that remaining pockets of wild-type individuals will experience a geometrical disadvantage after initial establishment of the drive. Importantly, this also needs to include any areas connected to the target area by moderate migration. The second strategy is to release the drives into an area with a higher carrying capacity than the surrounding area. This could allow a medium-threshold system to persist, and potentially even a high-threshold system if the difference in carrying capacity from the surrounding area is sufficiently large, with a sharp boundary. However, even a narrow migration corridor of similar density would likely result in the replacement of any high-threshold system. A 2L2T system could support a wider range of release strategies, but it should be expected to be invasive unless released into a strongly isolated population such as an island.

Our study marks a first step towards understanding the behavior of underdominance systems in realistic spatial populations, but more work remains to be done. While we restricted our analysis to only those underdominance systems that have been experimentally demonstrated, other designs have been proposed^33–38^ and could exhibit interesting dynamics that may change in spatially-continuous populations. *Wolbachia* population dynamics would also be interesting to model in comparable settings, and several studies have already shown that they can exhibit qualitatively similar dynamics to those we saw in low-threshold underdominance drives^19–28^. The results we presented in this study were all based on simulations, yet we hope that future studies will seek to develop an analytical understanding of the processes and parameters involved. The extension of Fisher and Bartonian wave models to underdominance systems could provide a fruitful avenue in this context^15,39–41^. Overall, our results clearly indicate that extensive modeling under realistic conditions must be performed to assess the chances of success for any underdominance application prior to its release.

## ACKNOWLEDGEMENTS

We thank Benjamin Haller for helpful discussions and assistance with the implementation of simulations models in SLiM. This study was supported by funding from New Zealand’s Predator Free 2050 program under award SS/05/01 to P.W.M, and grants from the National Institutes of Health under awards R01GM127418 to P.W.M, R21AI130635 to J.C., A.G.C. and P.W.M, and F32AI138476 to J.C.

---

**Figure S1.**
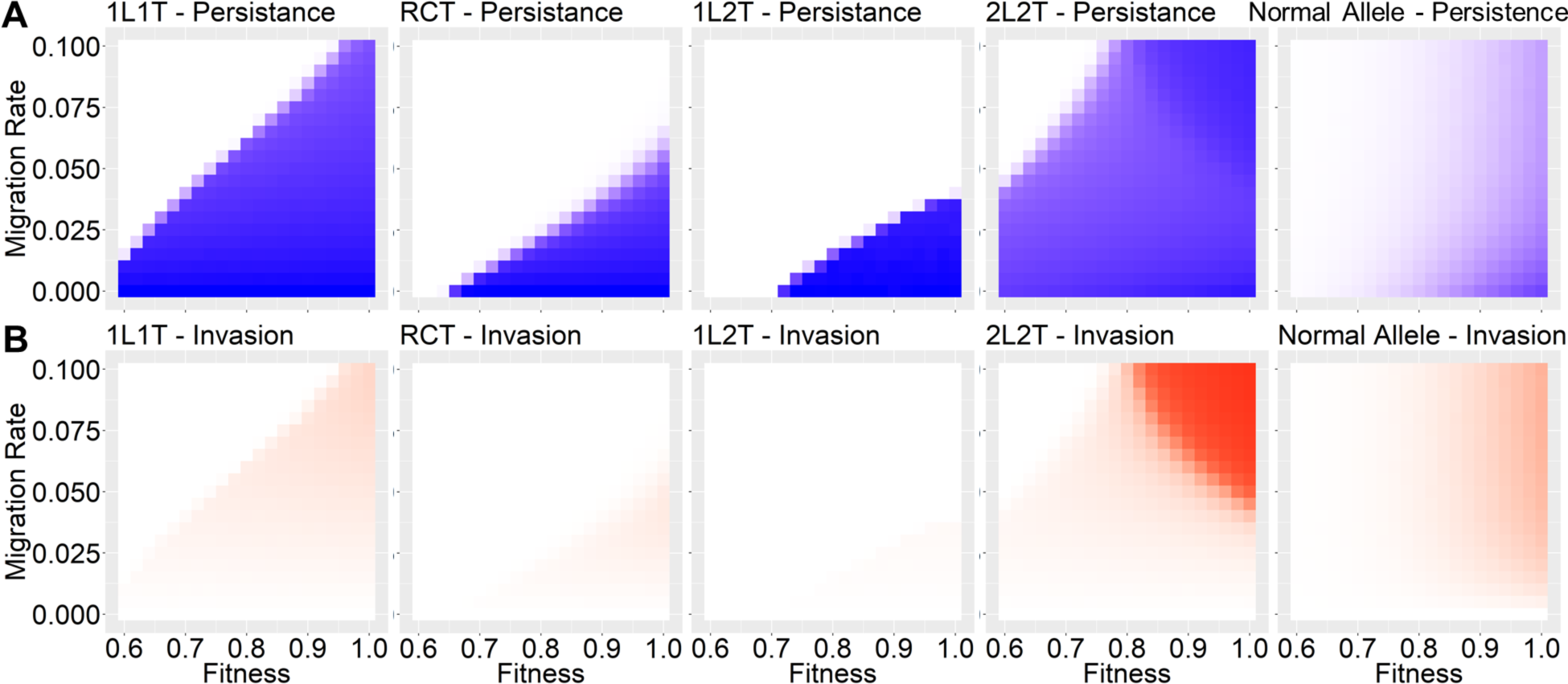
Panmictic release scenario. Heatmaps for all drives and a normal allele with the same scale and scenario as Figure 3 for (A) persistence and (B) invasion.

**Figure S2.**
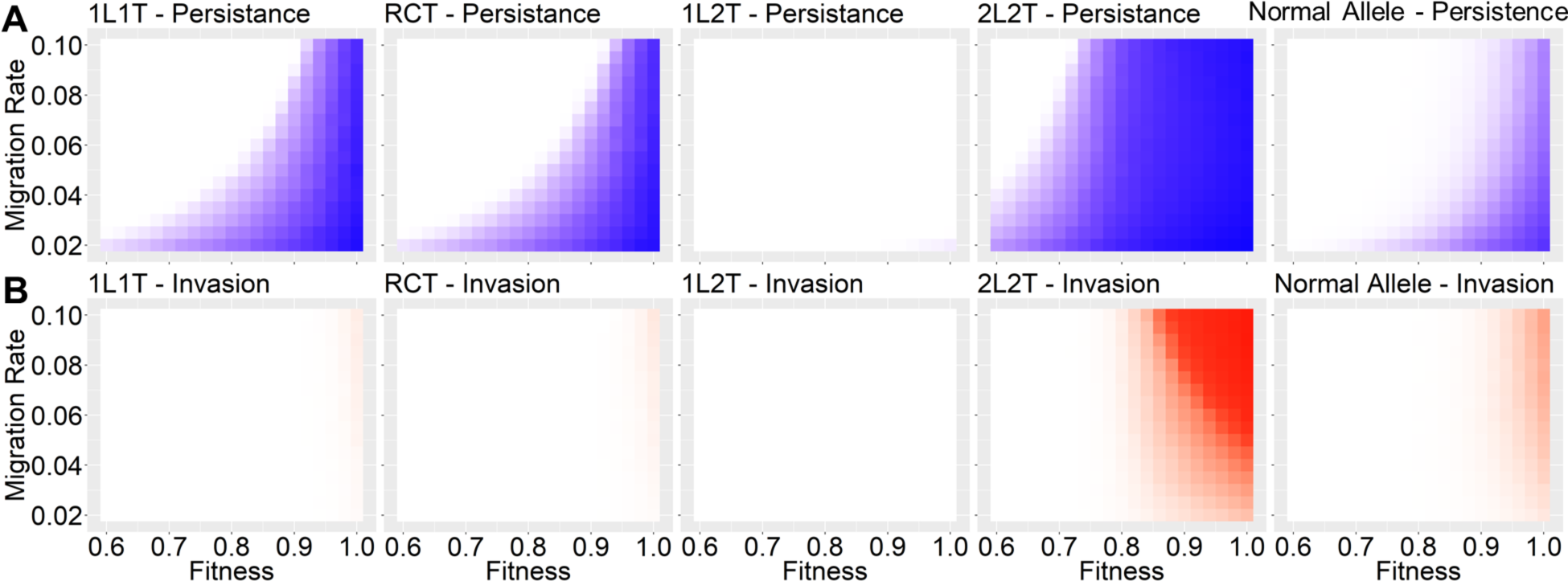
Linear scenario. Heatmaps for all drives and a normal allele with the same scale and scenario as Figure 4 for (A) persistence and (B) invasion.

**Figure S3.**
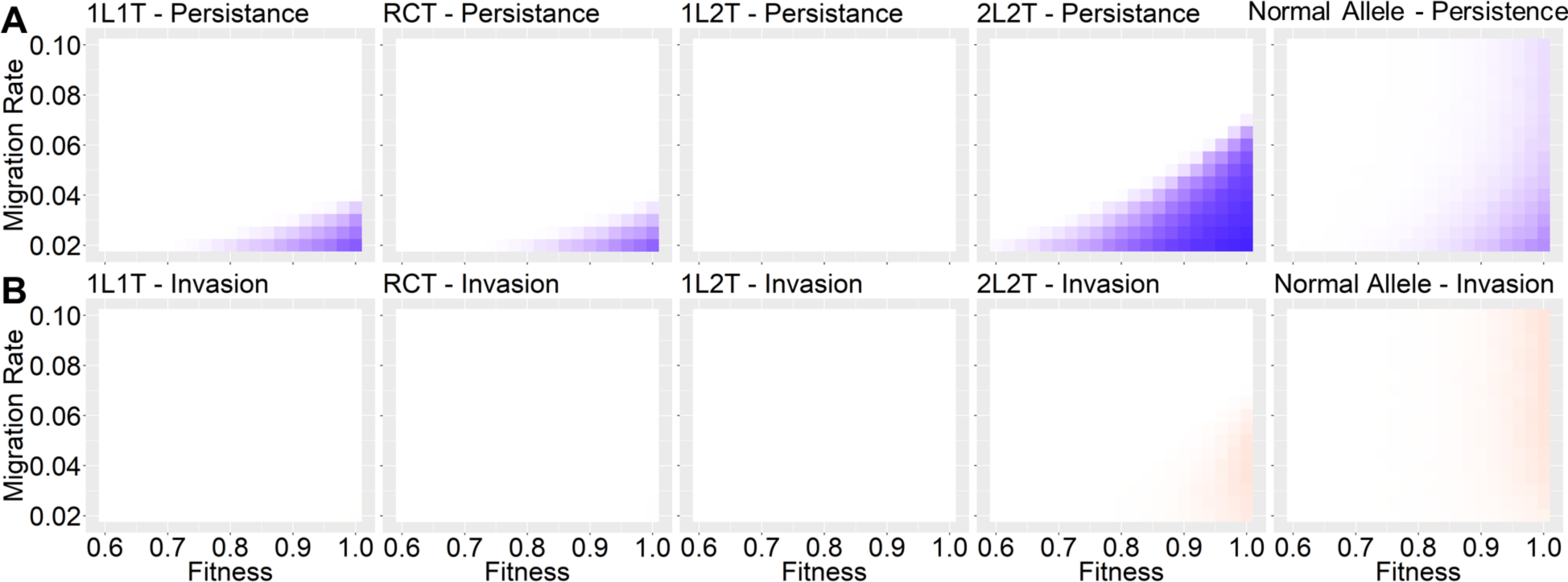
Circle release scenario. Heatmaps for all drives and a normal allele with the same scale and scenario as Figure 5 for (A) persistence and (B) invasion.

**Figure S4.**
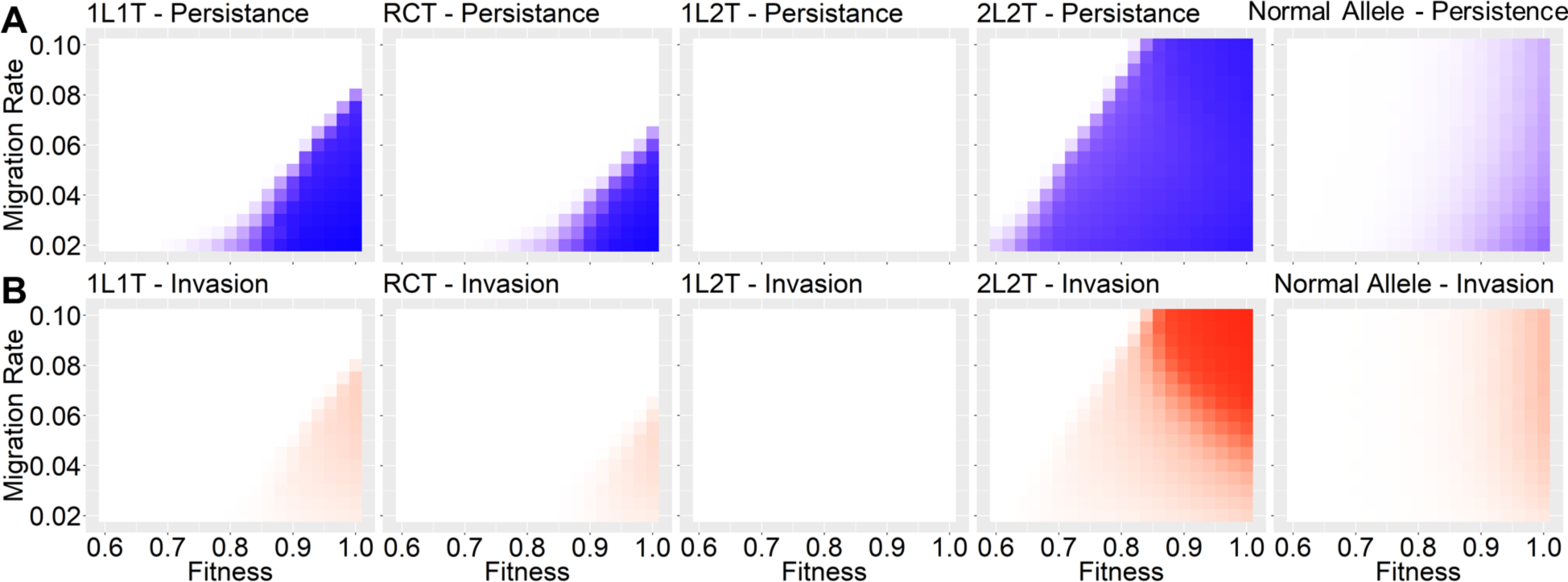
High-density circle release scenario, variable drive homozygote fitness and migration rate. Heatmaps for all drives and a normal allele with the same scale and scenario as Figure 6 for (A) persistence and (B) invasion.

**Figure S5.**
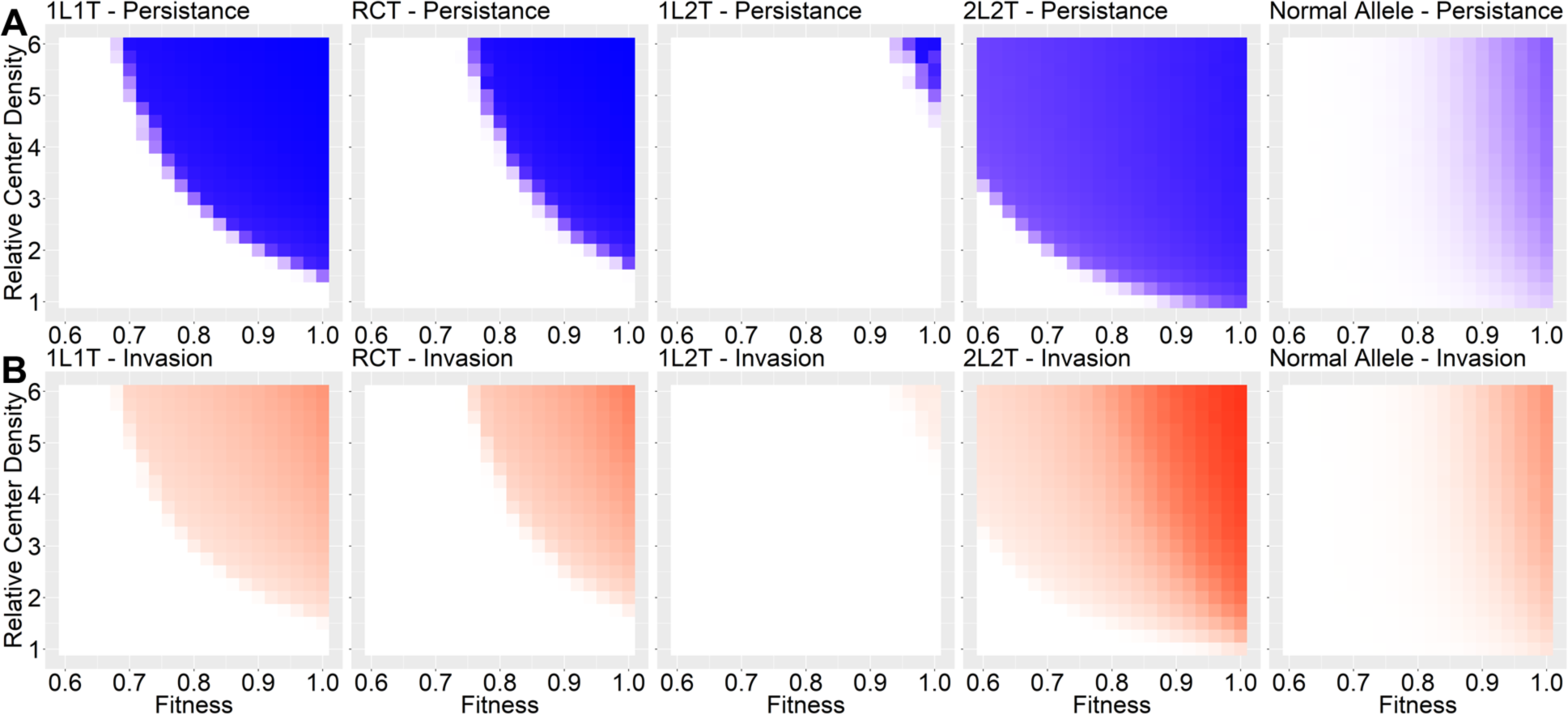
High-density circle release scenario, variable drive homozygote fitness and relative central density. Heatmaps for all drives and a normal allele with the same scale and scenario as Figure 6 for (A) persistence and (B) invasion.

**Figure S6.**
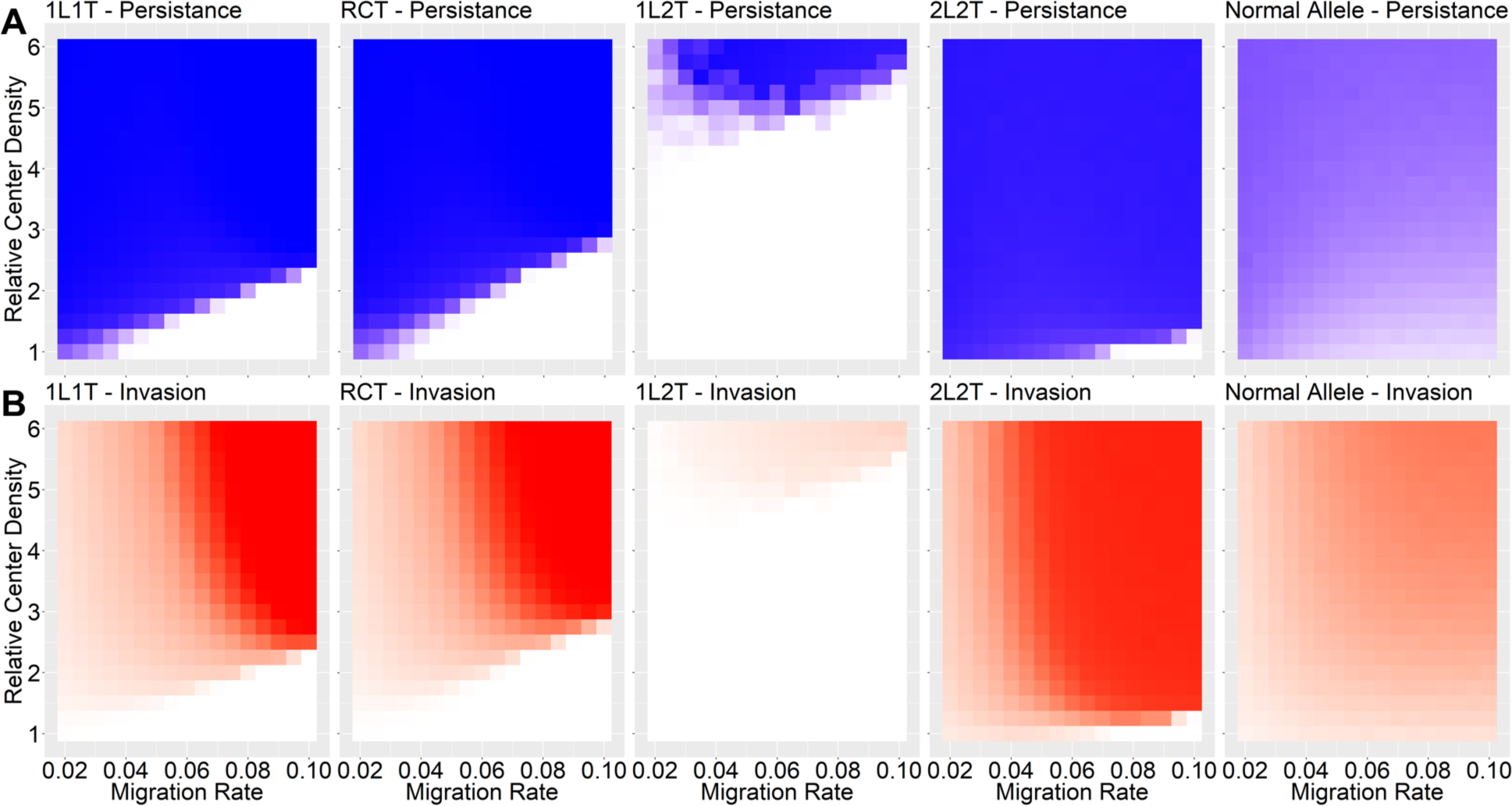
High-density circle release scenario, variable migration rate and relative central density. Heatmaps for all drives and a normal allele with the same scale and scenario as Figure 6 for (A) persistence and (B) invasion.

**Figure S7.**
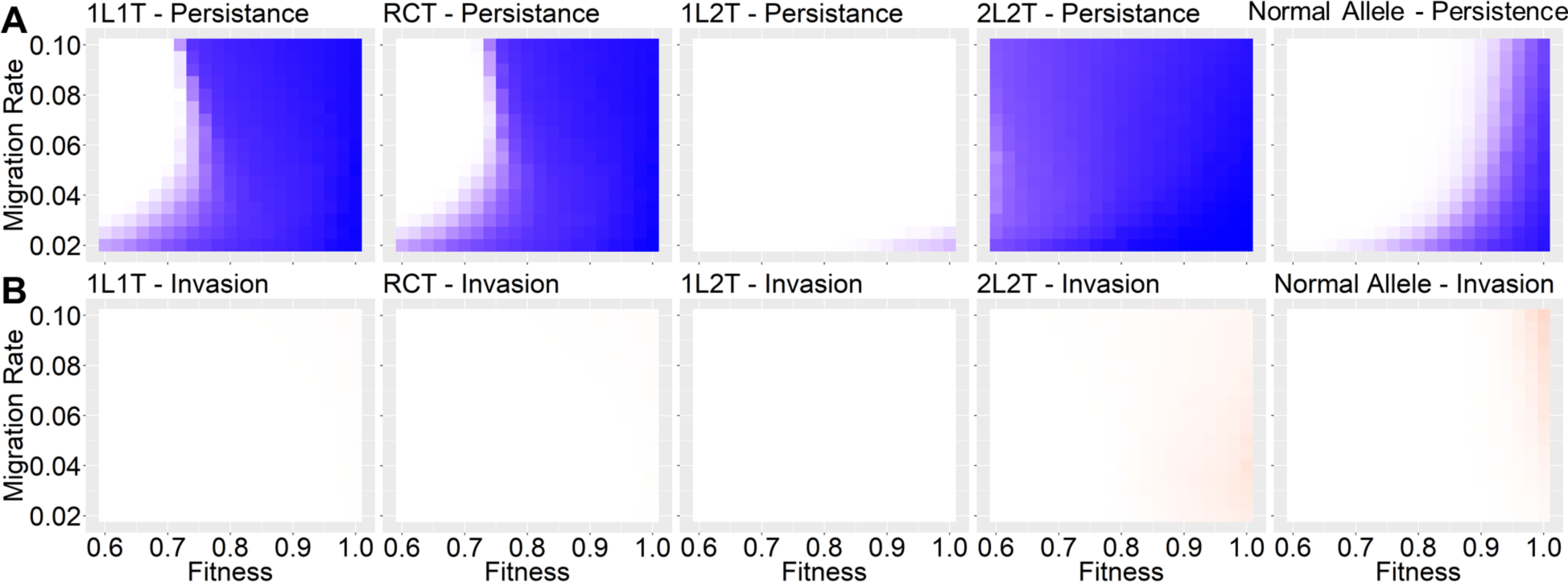
Migration corridor scenario, variable drive homozygote fitness and migration rate. Heatmaps for all drives and a normal allele with the same scale and scenario as Figure 7 for (A) persistence and (B) invasion.

**Figure S8.**
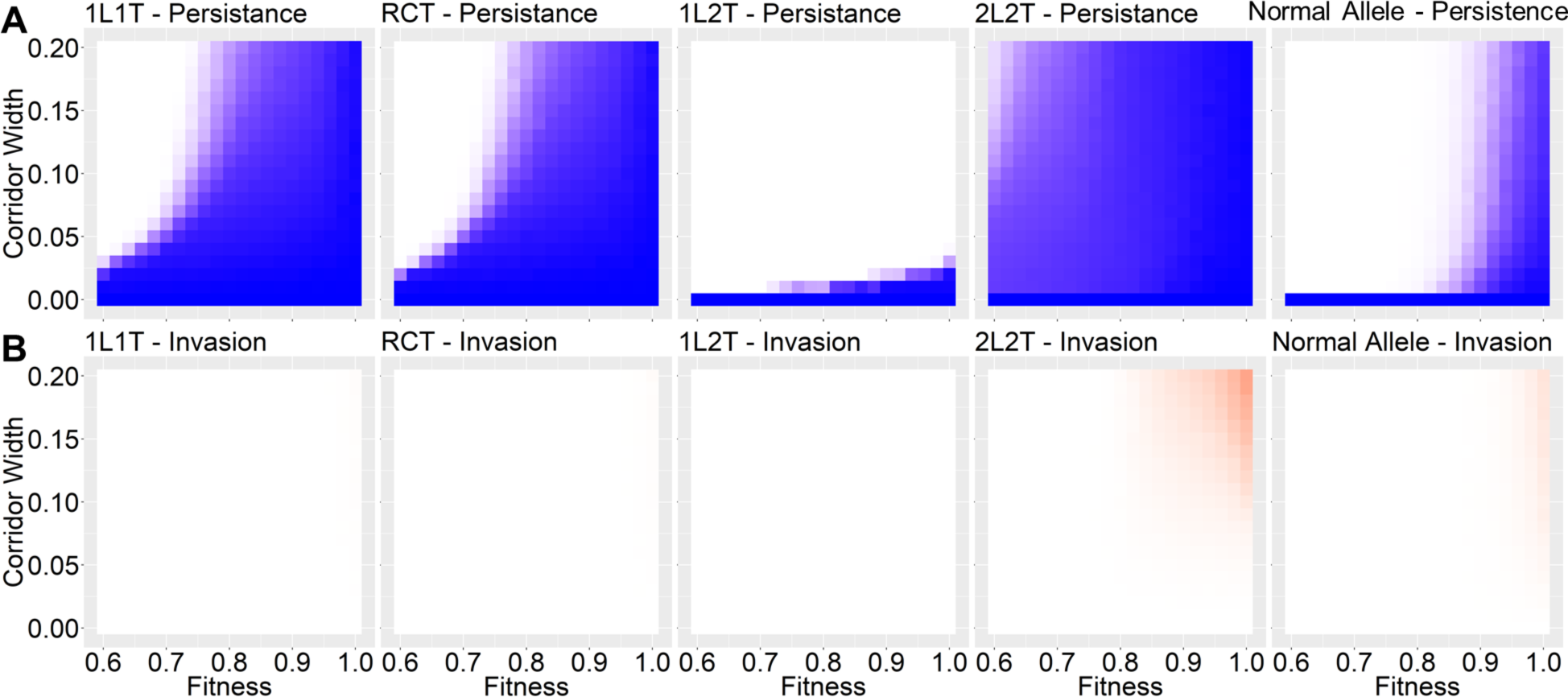
Migration corridor scenario, variable drive homozygote fitness and corridor width. Heatmaps for all drives and a normal allele with the same scale and scenario as Figure 7 for (A) persistence and (B) invasion.

**Figure S9.**
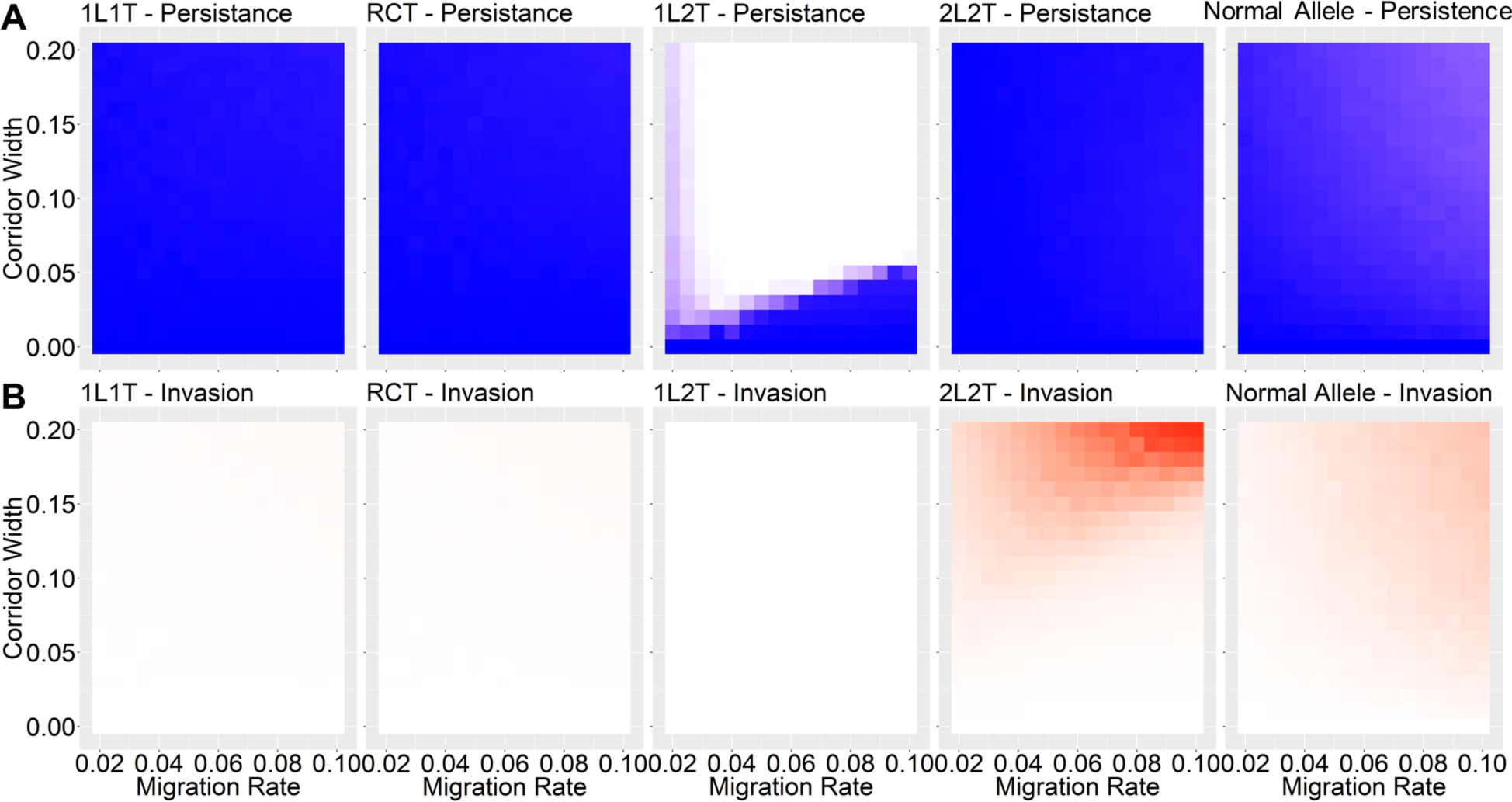
Migration corridor scenario, variable migration rate and corridor width. Heatmaps for all drives and a normal allele with the same scale and scenario as Figure 7 for (A) persistence and (B) invasion.

